# Mammalian RAVE couples V-ATPase assembly to organelle acidification and function

**DOI:** 10.64898/2025.12.10.693398

**Authors:** Nora S. Siefert, Andrea Zanotti, Anastasija Paneva, Martin Schneider, Dominic Helm, Wilhelm Palm

**Author notes:** **Corresponding author:** Correspondence to Wilhelm Palm. These authors contributed equally to this work (Nora S. Siefert, Andrea Zanotti).

## Abstract

Acidification of lysosomes, endosomes, and the Golgi underpins organelle-specific functions within the endomembrane system. This process is driven by vacuolar-type H⁺-ATPases (V-ATPases), which reversibly assemble from peripheral V₁ and membrane-integral V₀ domains to regulate organelle pH. In yeast, V₁–V₀ assembly at the vacuole is mediated by the RAVE complex, but V-ATPase assembly in mammalian cells remains less well understood. Here, we systematically define physiological roles of mammalian RAVE. Under basal conditions, mRAVE broadly promotes V-ATPase assembly and organelle acidification. Upon mTORC1 inactivation, mRAVE is recruited to lysosomes and required for the resulting increase in V-ATPase assembly and catabolic activity. Loss of mRAVE disrupts organelle acidification, leading to suppression of lysosomal catabolism, accumulation of dysfunctional lysosomes and compensatory lysosomal exocytosis. Restoring lysosomal pH rescues basal function in mRAVE-deficient cells but not the mTORC1-regulated increase in catabolic activity. Thus, mRAVE is an essential V-ATPase assembly factor that couples acidification to organelle function and nutrient signaling.

## Introduction

Luminal acidification of organelles is a central organizing principle through which eukaryotic cells establish functionally distinct compartments (Freeman *et al*, 2023). The most acidic compartment in mammalian cells is the lysosome, a catabolic organelle that contains numerous hydrolytic enzymes specialized for macromolecular breakdown (Ballabio & Bonifacino, 2020; Settembre & Perera, 2024). Several other organelles of the endomembrane system, including the Golgi apparatus and various endosomal compartments, also maintain a distinct acidic pH. These differences in luminal pH enable organelle-specific biochemical reactions and support diverse subcellular processes, such as membrane trafficking, receptor-ligand dissociation and signal transduction (Freeman *et al*., 2023). Consequently, perturbations in organelle acidification are linked to a range of human diseases, including cancer, neurodegeneration and genetic disorders, as well as aging (Eaton *et al*, 2021; Nixon & Rubinsztein, 2024; Tan & Finkel, 2023; Webb *et al*, 2021).

The acidification of organelles is mediated by vacuolar-type H⁺-ATPases (V-ATPases), multi-subunit proton pumps that use ATP hydrolysis to drive proton translocation across membranes (Collins & Forgac, 2020; Eaton *et al*., 2021; Vasanthakumar & Rubinstein, 2020). V-ATPases consist of two subcomplexes: a peripheral V_1_ domain, which hydrolyzes ATP, and a membrane-integral V_o_ domain, which translocates protons into the organelle lumen. The V_1_ and V_o_ domains exist in an equilibrium between separate complexes and the assembled V-ATPase. Regulation of V_1_ – V_o_ assembly constitutes a conserved mechanism that enables cells to dynamically adjust the number of active proton pumps (Kane, 1995; McGuire & Forgac, 2018; Ratto *et al*, 2022; Sautin *et al*, 2005; Sumner *et al*, 1995; Trombetta *et al*, 2003). The V_o_ domain is targeted to several organelles, which is mediated by different paralogs of V_o_ subunit a (Voa): Voa1 to endosomes and lysosomes, Voa2 to the Golgi and Voa3 (Tcirg1) to lysosomes. In specialized cell types, Voa subunits also direct the V_o_ domain to unique compartments, such as synaptic vesicles in neurons or the plasma membrane in osteoclasts and macrophages (Collins & Forgac, 2020; Vasanthakumar & Rubinstein, 2020). The regulation of V-ATPase subunit composition, subcellular localization and V_1_ – V_o_ domain assembly appears to be important to ensure that organelles maintain the luminal pH required for their specific function.

V-ATPase activity is tightly linked to lysosome function, establishing the acidic environment where the organelle’s catabolic enzymes are active (Ballabio & Bonifacino, 2020; Settembre & Perera, 2024). Lysosomes also integrate inputs from nutrient levels and growth factor signals to activate the kinase mechanistic target of rapamycin complex 1 (mTORC1), a central regulator of cellular metabolism and growth (Palm *et al*, 2015; Saxton & Sabatini, 2017). When mTORC1 is active, a substantial fraction of lysosomal V-ATPases is disassembled; conversely, nutrient starvation and resulting mTORC1 inactivation triggers V_1_ – V_o_ assembly, thereby rapidly increasing lysosomal acidification and enzyme activity (Palm *et al*., 2015; Ratto *et al*., 2022). Prolonged mTORC1 inactivation further promotes the expression of V-ATPase subunits via the transcription factor EB (TFEB) (Peña-Llopis *et al*, 2011). Together, these mechanisms couple nutrient signaling to acute and long-term adaptation of lysosomal catabolism.

Association of the V_1_ and V_o_ domains into a functional proton pump depends on dedicated assembly factors. In yeast, V-ATPase assembly is mediated by the RAVE (Regulator of H⁺-ATPase of Vacuolar and Endosomal membranes) complex, which consists of the central component Rav1 together with Rav2 and Skp1 (Jaskolka *et al*, 2021b; Seol *et al*, 2001). RAVE recruits cytosolic V_1_ domains to V_o_ domains at the vacuolar membrane and catalyzes their reassembly in response to glucose stimulation (Jaskolka *et al*, 2021a; Smardon *et al*, 2002). Mammals express two Rav1 homologs, Dmxl1 and Dmxl2, which associate with Wdr7—an unrelated protein not found in yeast RAVE (Jaskolka *et al*., 2021b; Kawabe *et al*, 2003). Early evidence implicating these proteins in V-ATPase assembly showed that Dmxl1, Dmxl2 and Wdr7 co-purify with the V-ATPase from mouse kidney and are required for reacidification of neutralized lysosomes (Merkulova *et al*, 2015). Consistently, co-depletion of Dmxl1 and Dmxl2 in cultured fibroblasts or depletion of Dmxl1 in mouse kidney suppresses V-ATPase assembly (Eaton *et al*, 2024; Ratto *et al*., 2022). Recent biochemical and structural studies identified Rogdi as the mammalian homolog of yeast Rav2 and defined a RAVE-related complex composed of Dmxl1 or Dmxl2, Wdr7 and Rogdi, referred to as metazoan RAVE (mRAVE) (Lee *et al*, 2025; Nardone *et al*, 2025; Winkley & Kane, 2025). mRAVE is recruited to damaged lysosomes with elevated pH and in turn promotes V-ATPase assembly to restore luminal acidification (Lee *et al*., 2025; Nardone *et al*., 2025). Whether mRAVE regulates V-ATPase assembly under basal conditions or in response to other physiological cues, however, has remained unclear. Moreover, beyond the lysosomal stress response, the role of mRAVE in supporting organelle functions has not been studied.

Here, we systematically define the role of mRAVE in organelle acidification and function. We show that mRAVE is broadly required for V-ATPase assembly across the endomembrane system under basal conditions and at lysosomes in response to mTORC1 signaling. Loss of mRAVE disrupts the acidification of various organelles and suppresses lysosomal catabolism, leading to accumulation of dysfunctional lysosomes and lysosomal exocytosis. These findings establish mRAVE as a general and essential V-ATPase assembly factor that safeguards basal and adaptive organelle function.

## Results

### mRAVE is required for basal assembly of the lysosomal V-ATPase

To systematically define mRAVE functions in mammalian cells, we generated inducible CRISPR/Cas9 knockouts (iKO) for each subunit—Wdr7, Dmxl1/2, and Rogdi—in mouse embryonic fibroblasts (MEFs). Cells deficient for Wdr7 or double-deficient for Dmxl1 and Dmxl2 proliferated at strongly reduced rates (Fig. 1A, B). Similar proliferation defects were observed across several mouse and human cell lines lacking Wdr7 (Fig. 1C), and these defects were rescued by expression of an sgRNA-resistant Wdr7 cDNA (Fig. EV1A, B). Loss of Rogdi had no overt effect (Fig. EV1C). Thus, Wdr7 and Dmxl1/2 are essential mammalian genes.

**Figure 1.**
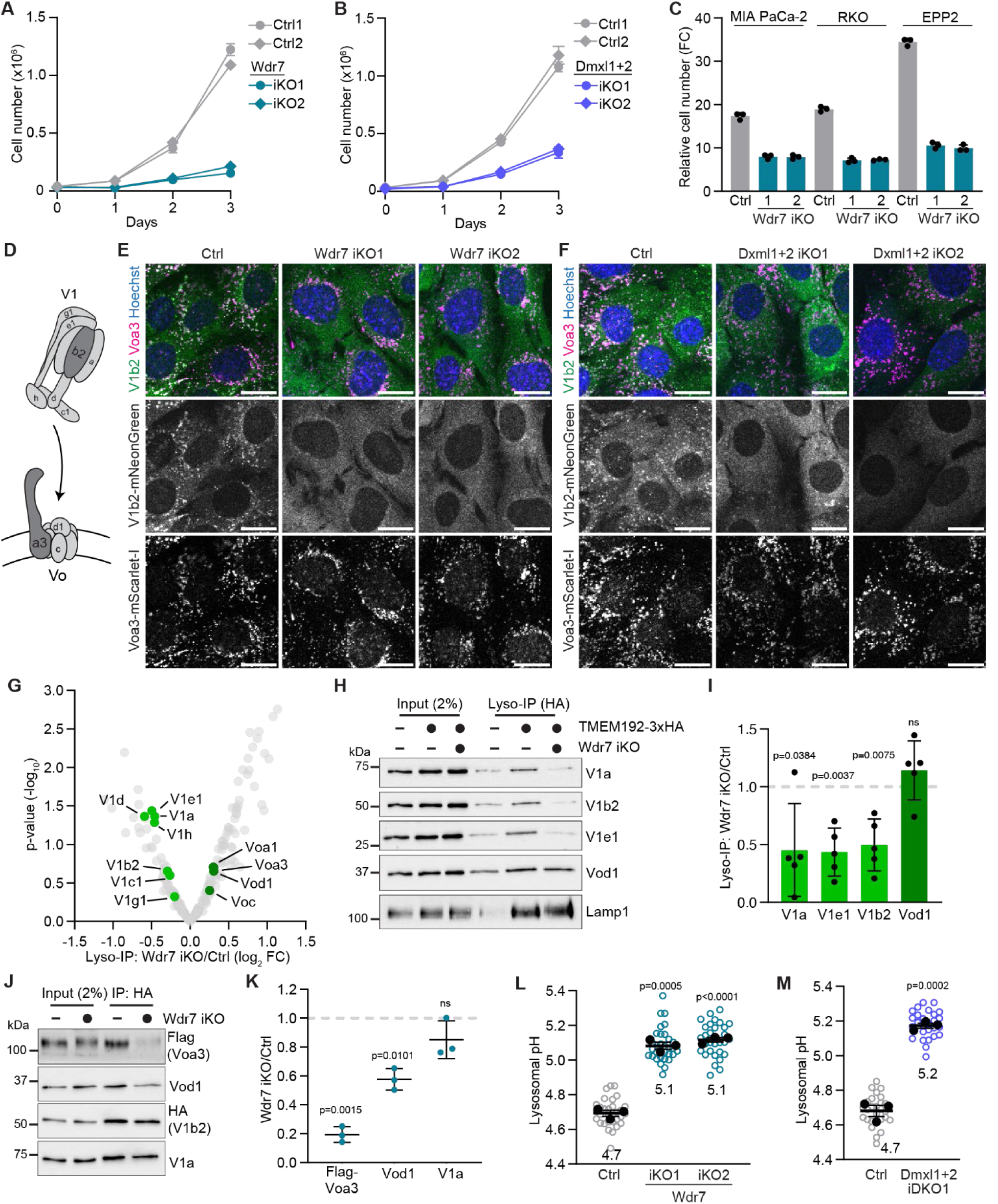
The mRAVE subunits Dmxl1/2 and Wdr7 are required for lysosomal V-ATPase assembly and acidification. **(A, B)** Proliferation of Wdr7 iKO (A) and Dmxl1 + Dmxl2 iKO (B) MEFs. **(C)** Fold change (FC) in cell number of indicated Wdr7 iKO cell lines after 3 days of proliferation. **(D)** Schematic of V-ATPase V_1_ – V_o_ domain assembly. **(E, F)** Subcellular localization of V1b2-mNeonGreen and Voa3-mScarlet-I in Wdr7 iKO (E) and Dmxl1 + Dmxl2 iKO (F) MEFs, analyzed by live-cell imaging. **(G)** Changes in lysosomal proteins in Wdr7 iKO MEFs, analyzed by Lyso-IP and label-free LC–MS. **(H)** Changes in V-ATPase subunit abundance in Lyso-IPs from Wdr7 iKO MEFs, analyzed by immunoblotting. **(I)** Quantification of V-ATPase subunit abundance in Lyso-IP immunoblots as in (H). **(J)** HA co-IP in Wdr7 iKO MEFs expressing HA-V1b2 and Flag-Voa3, analyzed by immunoblotting for V-ATPase subunits. **(K)** Quantification of V-ATPase subunit abundance in co-IP immunoblots as in (J). **(L, M)** Lysosomal pH in Wdr7 iKO (L) and Dmxl1 + Dmxl2 iKO (M) MEFs, quantified by ratiometric imaging of 10 kDa Dextran Oregon Green and Alexa Fluor 647. (**A – C**) Data are mean ± SD (n = 3 technical replicates). (**G**) n = 4 independent experiments. (**I, K**) Data are mean ± SD (n = 5 (I), n = 3 (K) independent experiments). (**L, M**) Data are replicate mean ± SEM (closed circles; n = 3 independent experiments) and fields of view (open circles; ≥ 10 per replicate). *p* values were calculated using a two-tailed unpaired t-test with Welch correction (I and K) or a two-tailed unpaired t-test (L and M). Scale bars, 20 µm.

To determine whether mRAVE is required for V-ATPase assembly at lysosomes, we labeled the peripheral V1 domain and membrane-integral V_o_ domain by stable expression of V1b2-mNeonGreen and Voa3-mScarlet-I, respectively. In control cells, V1b2 partially co-localized with Voa3 but was also diffusely present throughout the cytosol, reflecting the co-occurrence of assembled and disassembled V-ATPase pools (Fig. 1D–F) (Ratto *et al*., 2022). In cells deficient for Wdr7 or double-deficient for Dmxl1 and Dmxl2, however, most V_1_ domains localized to the cytosol, with only a minor fraction localizing to Voa3-containing organelles. To examine endogenous V-ATPase subunits, we enriched lysosomes by immunoprecipitating the lysosomal membrane protein TMEM192-HA (Lyso-IP (Abu-Remaileh *et al*, 2017)) and analyzed the resulting fractions by immunoblotting as well as liquid chromatography–mass spectrometry (LC–MS) and label-free quantification (LFQ). Lyso-IP efficiently enriched lysosomal proteins, including V-ATPase subunits, while depleting proteins that localize to non-lysosomal compartments, confirming high purity of lysosomal fractions (Fig. EV1D–F; Dataset EV1). The abundance of V_1_ domain subunits was significantly decreased in lysosomal fractions from Wdr7-deficient cells (Fig. 1G–I; Dataset EV2). In contrast, V_o_ domain subunits were not decreased. To examine the physical association between V_1_ and V_o_ domains directly, we conducted co-immunoprecipitation (co-IP) experiments using HA-V1b2 as bait. V_o_ domain subunits were significantly reduced in HA-V1b2 immunoprecipitates from Wdr7-deficient cells, whereas V_1_ domain subunit interactions remained intact (Fig. 1J, K). To determine the functional consequences of disrupted V_1_ – V_o_ domain assembly, we quantified lysosomal pH by ratiometric imaging of lysosomally loaded dextran Oregon Green, whose fluorescence is quenched at acidic pH, and pH-insensitive dextran Alexa Fluor 647. Lysosomal pH increased from ∼4.7 to 5.1–5.2 in cells deficient for Wdr7 or double-deficient for Dmxl1 and Dmxl2, consistent with the decrease of active V-ATPases (Fig. 1L, M). Thus, mRAVE is critical for basal assembly of lysosomal V-ATPases and, consequently, for lysosomal acidification.

To examine the contributions of the two Dmxl homologs, Dmxl1 and Dmxl2, we depleted each protein individually. Loss of either Dmxl1 or Dmxl2 reduced the interaction between V_1_ domains and Voa3-containing V_o_ domains (Fig. EV2A–D), shifted the localization of V1b2-mNeonGreen toward the cytosol (Fig. EV2E), elevated lysosomal pH (Fig. EV2F) and compromised cell proliferation (Fig. EV2G). Thus, cells deficient for either Dmxl1 or Dmxl2 displayed similar, albeit less pronounced, phenotypes to cells deficient for both homologs, suggesting partially non-redundant functions. Given the penetrant phenotypes observed in Wdr7-deficient cells, we focused subsequent analyses on Wdr7 as an essential mRAVE subunit.

### mRAVE is a general assembly factor for V-ATPase complexes

To examine whether mRAVE is required for assembly of distinct V-ATPase complexes at different subcellular locations, we expressed fluorescently labeled homologs of the Voa subunit: Voa1, which localizes to the endo-lysosomal compartment, including Voa3-positive lysosomes, and Voa2, which localizes to the Golgi (Fig. EV3A, B) (Collins & Forgac, 2020; Vasanthakumar & Rubinstein, 2020). In control cells, V1b2-mNeonGreen showed robust co-localization with Voa1-mScarlet-I and Voa2-mScarlet-I (Fig. 2A, B). In Wdr7-deficient cells, however, very little co-localization of V1b2 with either Voa1- or Voa2-containing organelles was detectable. Consistently, Flag-Voa1 and Flag-Voa2 co-immunoprecipitated less with V1b2 in Wdr7-deficient cells (Fig. 2C–F). To assess the functional impact of decreased V-ATPase assembly at the Golgi, we measured luminal pH using the Golgi-specific pH sensor MGAT-SEP-mRuby (Liu *et al*, 2023). Golgi pH increased from 6.4 in control cells to 7.3 in Wdr7-deficient cells, indicating near-complete neutralization of the organelle lumen (Fig. 2G). We were unable to quantify endosomal pH due to the lack of compartment-specific pH sensors. Together, these results establish mRAVE as a general assembly factor for V-ATPase complexes at different organelles.

**Figure 2.**
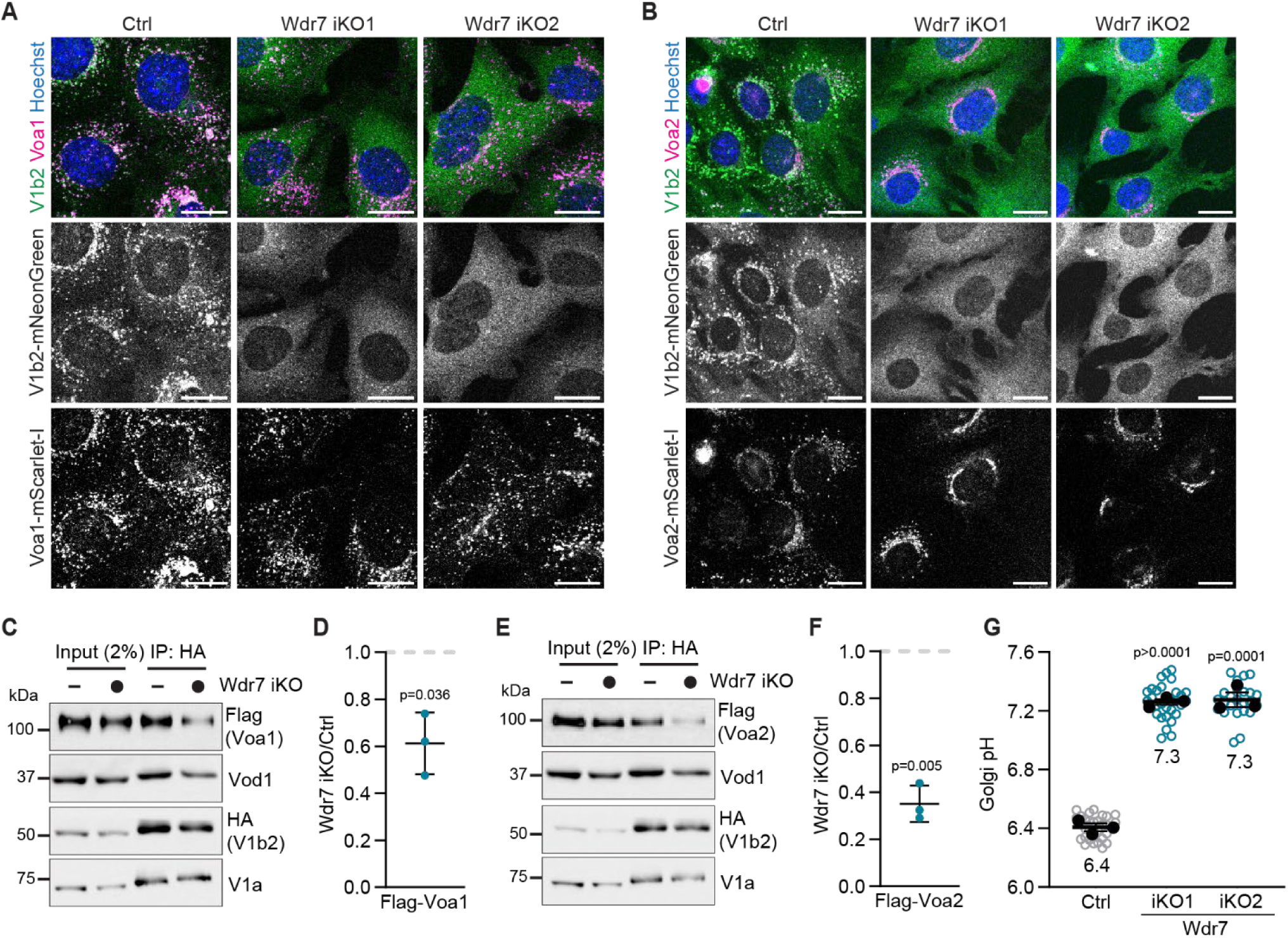
mRAVE supports V-ATPase assembly at different organelles. **(A**, **B)** Subcellular localization of V1b2-mNeonGreen and Voa1-mScarlet-I (A) or Voa2-mScarlet-I (B) in Wdr7 iKO MEFs, analyzed by live-cell imaging. **(C)** HA co-IP in Wdr7 iKO MEFs expressing HA-V1b2 and Flag-Voa1, analyzed by immunoblotting for V-ATPase subunits. **(D)** Quantification of V-ATPase subunit abundance in co-IP immunoblots as in (C). **(E)** HA co-IP in Wdr7 iKO MEFs expressing HA-V1b2 and Flag-Voa2, analyzed by immunoblotting for V-ATPase subunits. **(F)** Quantification of V-ATPase subunit abundance in co-IP immunoblots as in (E). **(G)** Golgi pH in Wdr7 iKO MEFs, quantified by ratiometric imaging of MGAT-SEP-mRuby. (**D, F**) Data are mean ± SD (n = 3 independent experiments). (**G**) Data are replicate mean ± SEM (closed circles; n = 3 independent experiments) and fields of view (open circles; ≥ 9 per replicate). *p* values were calculated using a two-tailed unpaired t-test with Welch correction (**D, F)** or a two-tailed unpaired t-test (G). Scale bars, 20 µm.

### mRAVE promotes assembly of lysosomal V-ATPases in response to mTORC1 inhibition

To further define the role of mRAVE in V-ATPase assembly, we identified V_1_ domain interaction partners by co-IP and LC–MS. V_1_ and V_o_ domain subunits were highly enriched in HA-V1b2 immunoprecipitates, confirming specificity of the co-IP (Fig. 3A; Dataset EV3). V1b2 immunoprecipitates were also enriched in subunits of the Ragulator complex (Lamtor1–4), which associates with lysosomal V-ATPases (Zoncu *et al*, 2011). Notably, V1b2 co-IPs robustly recovered the different mRAVE subunits, Dmxl1, Wdr7 and Rogdi. Immunoblotting confirmed that HA-V1b2 co-precipitated both V5-tagged Wdr7 and V_o_ domain subunits (Fig. 3B). Thus, the V_1_ domain interacts with both the V_o_ domain and mRAVE, consistent with previous results (Lee *et al*., 2025; Nardone *et al*., 2025).

**Figure 3.**
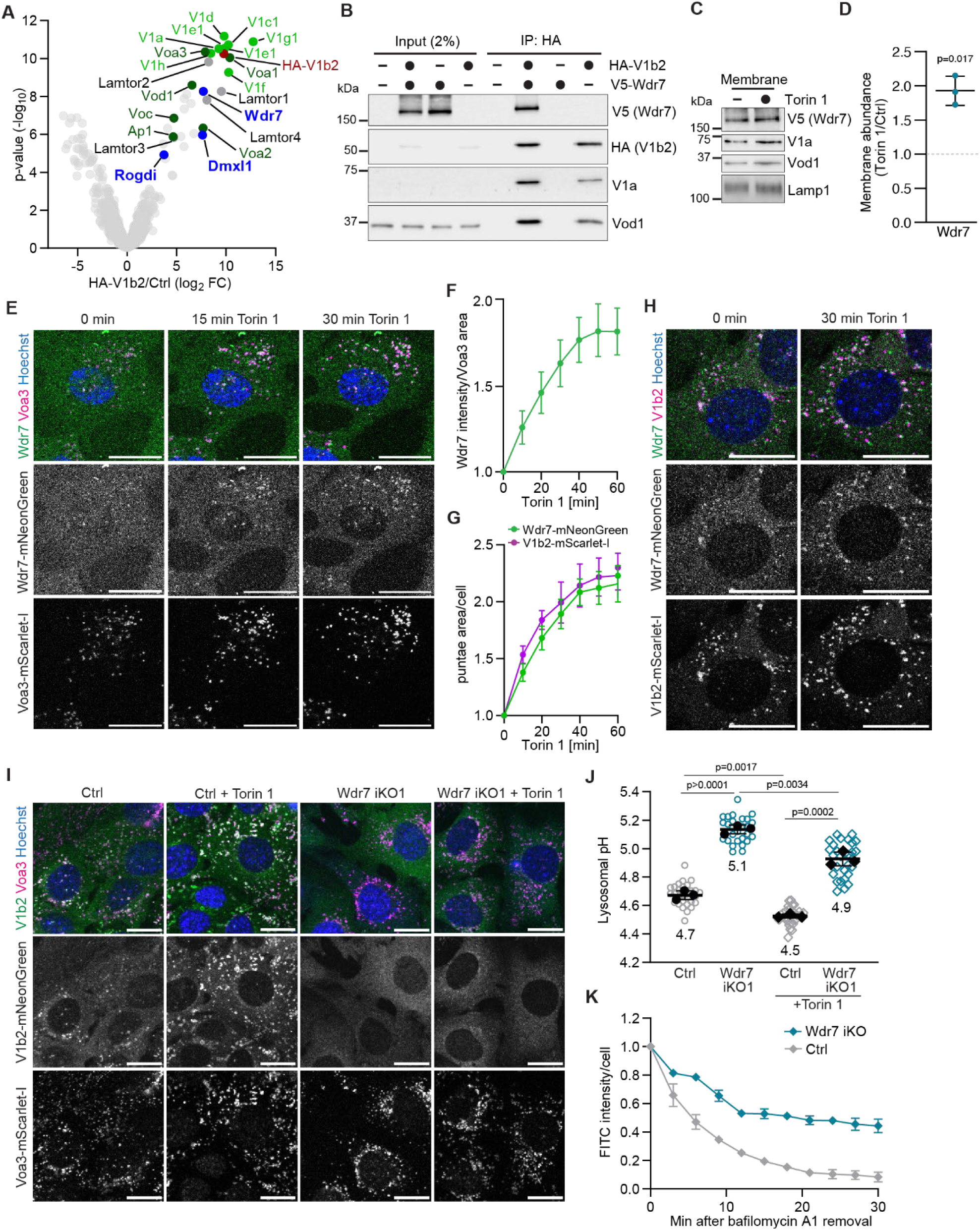
mRAVE couples mTORC1 signaling to V-ATPase assembly. **(A)** HA co-IP in MEFs expressing HA-V1b2, analyzed by label-free LC–MS. **(B)** HA co-IP in MEFs expressing HA-V1b2 and V5-Wdr7, analyzed by immunoblotting. **(C)** V5-Wdr7 abundance in membrane fractions of dextran-loaded lysosomes from MEFs after 1 h ± torin 1 [400 nM], analyzed by immunoblotting. **(D)** V5-Wdr7 abundance change in membrane fractions, quantified by immunoblotting as shown in (C). **(E)** Subcellular localization of Wdr7-mNeonGreen and Voa3-mScarlet-I in MEFs over time after addition of torin 1 [400 nM], analyzed by live-cell imaging. **(F)** Change of Wdr7-mNeonGreen intensity in Voa3-mScarlet-I-containing lysosomes, quantified by live-cell imaging as shown in (E). **(G)** Change in organellar Wdr7-mNeonGreen and V1b2-mScarlet-I, quantified by live-cell imaging as shown in (H). **(H)** Subcellular localization of Wdr7-mNeonGreen and V1b2-mScarlet-I in MEFs over time after addition of torin 1 [400 nM], analyzed by live-cell imaging. **(I)** Subcellular localization of V1b2-mNeonGreen and Voa3-mScarlet-I in Wdr7 iKO MEFs after 1 h ± torin 1 [400 nM], analyzed by live-cell imaging. **(J)** Lysosomal pH in Wdr7 iKO cells after 1 h ± torin 1 [400 nM], quantified by ratiometric imaging of 10 kDa Dextran Oregon Green and Alexa Fluor-647. **(K)** Fluorescence quenching of lysosomally loaded FITC-dextran in Wdr7 iKO MEFs after 1 h bafilomycin A1 [20 nM] and torin 1 [400 nM] and subsequent bafilomycin A1 removal. (**A**) n = 5 (HA-V1b2) and n = 4 (Ctrl) independent experiments. (**D**) Data are mean ± SD (n = 3 independent experiments). (**F, G**) Data are mean ± SD (≥ 5,000 organelles in ≥ 8 fields of view with a total of ≥ 70 cells). (**J**) Data are replicate mean ± SEM (closed circles; n = 3 independent experiments) and fields of view (open circles; ≥ 10 per replicate). (**K**) Data are mean ± SD (n = 3 technical replicates). *p* values were calculated using a two-tailed unpaired t-test with Welch correction (D) or a two-tailed unpaired t-test (J). Scale bars, 20 µm.

mTORC1 inactivation triggers the recruitment of V_1_ domains to lysosomal V_o_ domains to assemble active proton pumps (Ratto *et al*., 2022). To investigate a potential role of mRAVE in this process, we treated cells with the mTOR kinase inhibitor torin 1 and examined the impact on mRAVE localization. Torin 1 treatment increased the abundance of V5-Wdr7 in organelle fractions enriched for dextran-weighted lysosomes (Fig. 3C, D). Live-cell imaging showed that Wdr7-mNeonGreen was predominantly cytosolic under basal conditions but rapidly accumulated at Voa3-mScarlet-I-containing lysosomes in response to torin 1 (Fig. 3E, F). Consistently, we observed co-accumulation of Wdr7-mNeonGreen and V1b2-mScarlet-I at organelles over a comparable time period (Fig. 3G, H). This suggests that mRAVE and V_1_ domains are concertedly recruited to lysosomes upon mTORC1 inactivation.

Next, we examined whether mRAVE is required for mTORC1-regulated V-ATPase assembly. Torin 1 triggered recruitment of V1b2 to Voa3-containing lysosomes in control cells, indicating enhanced V-ATPase assembly (Fig. 3I) (Ratto *et al*., 2022). In Wdr7-deficient cells, however, V1b2 did not accumulate at lysosomes upon torin 1 treatment but rather remained cytosolic (Fig. 3I). Consequently, loss of Wdr7 caused a marked increase in lysosomal pH under both basal and torin 1-treated conditions (Fig. 3J). Thus, mRAVE supports the increase in lysosomal V-ATPase assembly and acidification in response to mTORC1 inactivation. To directly monitor V-ATPase-dependent acidification, we pre-loaded lysosomes with FITC-dextran, neutralized luminal pH with the V-ATPase inhibitor bafilomycin A1, and enhanced V-ATPase assembly with torin 1 (Ratto *et al*., 2022; Yoshimori *et al*, 1991). Bafilomycin A1 washout promptly restored V-ATPase activity in control cells, resulting in rapid FITC quenching (Fig. 3K). In contrast, the rate and extent of FITC quenching was strongly decreased in Wdr7-deficient cells. Notably, torin 1 did not affect Golgi pH, suggesting that V-ATPase assembly at the Golgi depends on mRAVE but is not regulated by mTORC1 signaling (Fig. EV3C). Together, these results indicate that mRAVE is critical for the enhanced lysosomal V-ATPase assembly and luminal acidification in response to mTORC1 inactivation.

### mRAVE deficiency causes accumulation of dysfunctional lysosomes

Because lysosomes in mRAVE-deficient cells were partially acidic—despite the strong reduction in assembled V-ATPases—we tested whether the remaining acidification depends on V-ATPase activity. Treating Wdr7-deficient cells with bafilomycin A1 further elevated lysosomal pH, indicating that residual V-ATPases sustained some luminal acidification (Fig. 4A). To assess the functional consequences of decreased acidification, we examined lysosome abundance by staining for Lamp2, a ubiquitous lysosomal membrane protein (Chen *et al*, 1985). Loss of Wdr7 caused a significant accumulation of Lamp2-positive organelles (Fig. 4B, C). This change was accompanied by a marked increase in lysotracker red, a fluorescent dye whose accumulation in cells reports on lysosomal abundance and acidity (Fig. 4D, E). Bafilomycin A1 treatment abolished lysotracker signal (Fig. EV4A, B) suggesting that without mRAVE, lysosomes remained sufficiently acidic to retain the dye. Together, these results show that loss of mRAVE leads to accumulation of lysosomes with diminished acidity.

**Figure 4.**
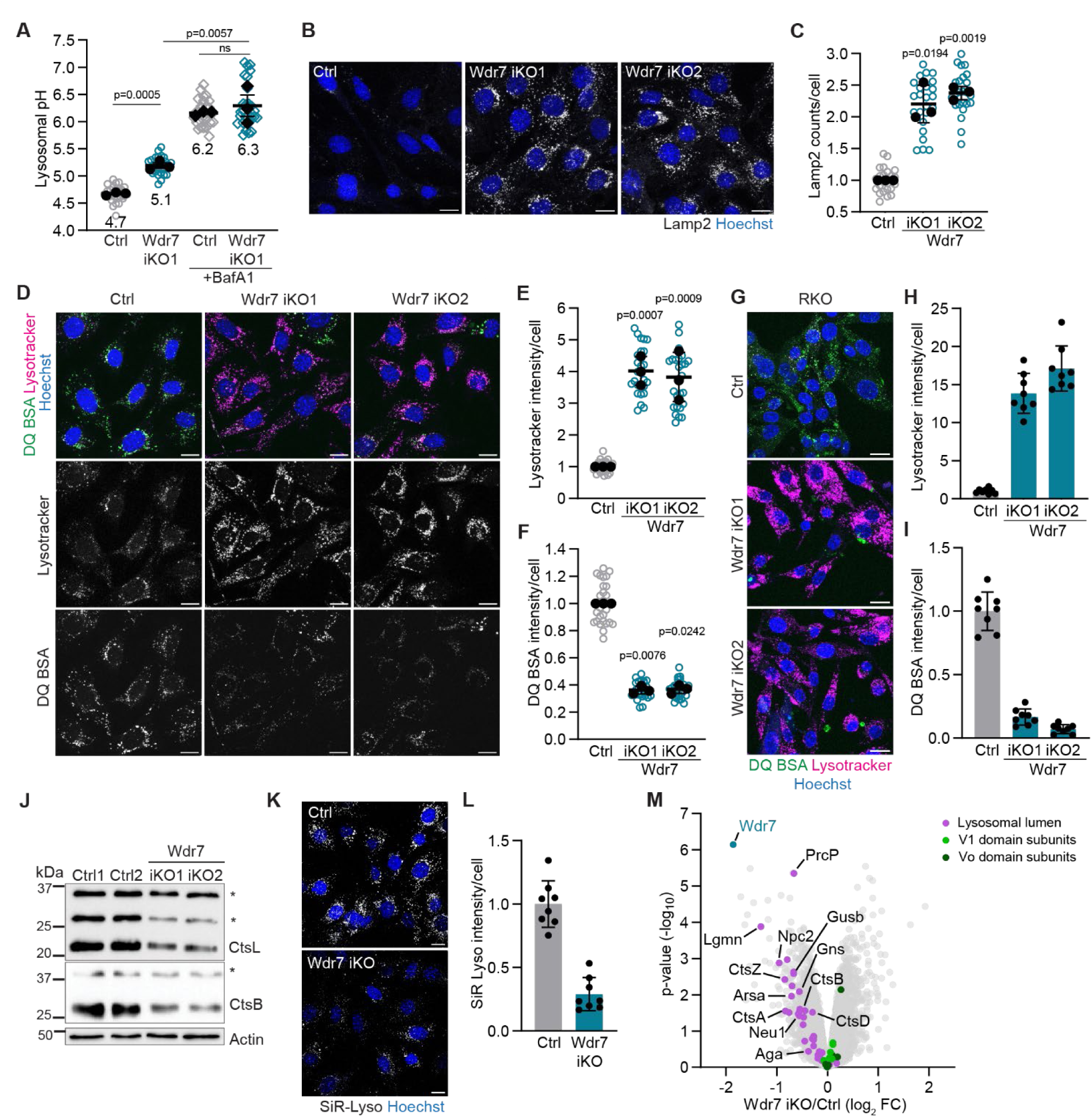
Loss of mRAVE causes accumulation of lysosomes with impaired catabolic activity. **(A)** Lysosomal pH in Wdr7 iKO MEFs after 1.5 h ± bafilomycin A1 [200 nM], quantified by ratiometric imaging of 10 kDa Dextran Oregon Green and Alexa Fluor 647. **(B)** Lysosome accumulation in Wdr7 iKO MEFs, analyzed by Lamp2 immunofluorescence (IF). **(C)** Quantification of Lamp2 accumulation in Wdr7 iKO MEFs as shown in (B). **(D)** Lysotracker accumulation and DQ BSA degradation in Wdr7 iKO MEFs, analyzed by live-cell imaging. **(E**, **F)** Quantification of Lysotracker (E) and DQ BSA (F) fluorescence in Wdr7 iKO MEFs as shown in (D). **(G)** Lysotracker accumulation and DQ BSA degradation in Wdr7 iKO RKO cells, analyzed by live-cell imaging. **(H, I)** Quantification of Lysotracker (H) and DQ BSA (I) fluorescence in Wdr7 iKO RKO cells as shown in (G). **(J)** Mature cathepsin abundance in Wdr7 iKO MEFs, analyzed by immunoblotting. Asterisk denotes immature proenzymes. **(K)** Active cathepsin D abundance in Wdr7 iKO MEFs, analyzed by SiR-Lysosome imaging. **(L)** Quantification of SiR-Lysosome fluorescence of cells shown in (K). **(M)** Changes in the proteome of Wdr7 iKO MEFs, quantified by label-free LC-MS. (**A, C, E, F**) Data are replicate mean ± SEM (closed circles; n = 3 independent replicates) and fields of view (open circles; ≥ 8 per replicate). (**H, I, L**) Data are mean ± SD and fields of view (n = 8). (**M**) n = 4 independent experiments. *p* values were calculated using a two-tailed unpaired t-test with Welch correction (C, E and F) or a two-tailed unpaired t-test (A). Scale bars, 20 µm.

To examine the impact of mRAVE on the catabolic activity of lysosomes, we incubated cells with DQ BSA, an albumin probe, whose fluorescence becomes dequenched upon degradation in lysosomes (Palm *et al*., 2015). Loss of Wdr7 led to a significant decrease in DQ BSA fluorescence dequenching in MEFs (Fig. 4D, F), as well as in several mouse and human cancer cell lines (Fig. 4G, I; Fig. EV4C-H). A comparable decrease in DQ BSA degradation was observed in MEFs double-deficient for Dmxl1 and Dmxl2 (Fig. EV4I, K). In each of these cell lines, this was accompanied by a strong accumulation of lysotracker-positive organelles (Fig. 4D, E, G, H; Fig. EV4C–H). We also examined the individual impact of Dmxl1, Dmxl2 and Rogdi. Loss of Dmxl1 or Dmxl2 moderately decreased DQ BSA degradation and increased lysotracker signal (Fig. EV5A–C), while loss of Rogdi had no effect (Fig. EV5D–F), mirroring the impact of these genes on cell proliferation.

To further characterize the defects in the lysosomal catabolism resulting from loss of mRAVE, we examined the abundance of lysosomal enzymes. Wdr7-deficient MEFs showed a marked reduction in the mature forms of cathepsins B and L, two ubiquitous lysosomal proteases (Fig. 4J) (Turk *et al*, 2012). Consistently, loss of Wdr7 led to a strong decrease in SiR-lysosome (Lukinavičius *et al*, 2016), a fluorescent probe that binds to active cathepsin D (Fig. 4K, L). To further characterize the phenotypes of mRAVE-deficient cells, we quantified cellular proteins in Wdr7-deficient MEFs by LC–MS and LFQ. Loss of Wdr7 led to an overall decrease in the abundance of lysosomal enzymes and other luminal lysosomal proteins (Fig. 4M; Dataset EV4). This decrease in lysosomal enzymes and lysosomal proteolytic activity in the absence of mRAVE, despite an expanded lysosomal compartment, indicates a strong defect in lysosomal catabolic activity.

### mRAVE deficiency results in lysosomal exocytosis

We noted that mRAVE-deficient cells appeared to have normal levels of immature pro-cathepsins, which are proteolytically processed into the mature enzymes after reaching the lysosome (Fig. 4J) (Braulke *et al*, 2024). Because this suggested perturbed trafficking of lysosomal proteins, we analyzed cellular supernatants by immunoblotting. Immature pro-cathepsins B and L were strongly increased in supernatants from cells deficient for Wdr7 or double-deficient for Dmxl1 and Dmxl2 (Fig. 5A, B). To more broadly examine lysosomal enzyme secretion, we quantified proteins in cellular supernatants using LC-MS and LFQ. Lysosomal enzymes were overall increased in supernatants from cells deficient for Wdr7 or double-deficient for Dmxl1 and Dmxl2, with little change in constitutively secreted proteins (Fig. 5C, D; Dataset EV5). Thus, loss of mRAVE leads to increased secretion of lysosomal enzymes into the extracellular space.

**Figure 5.**
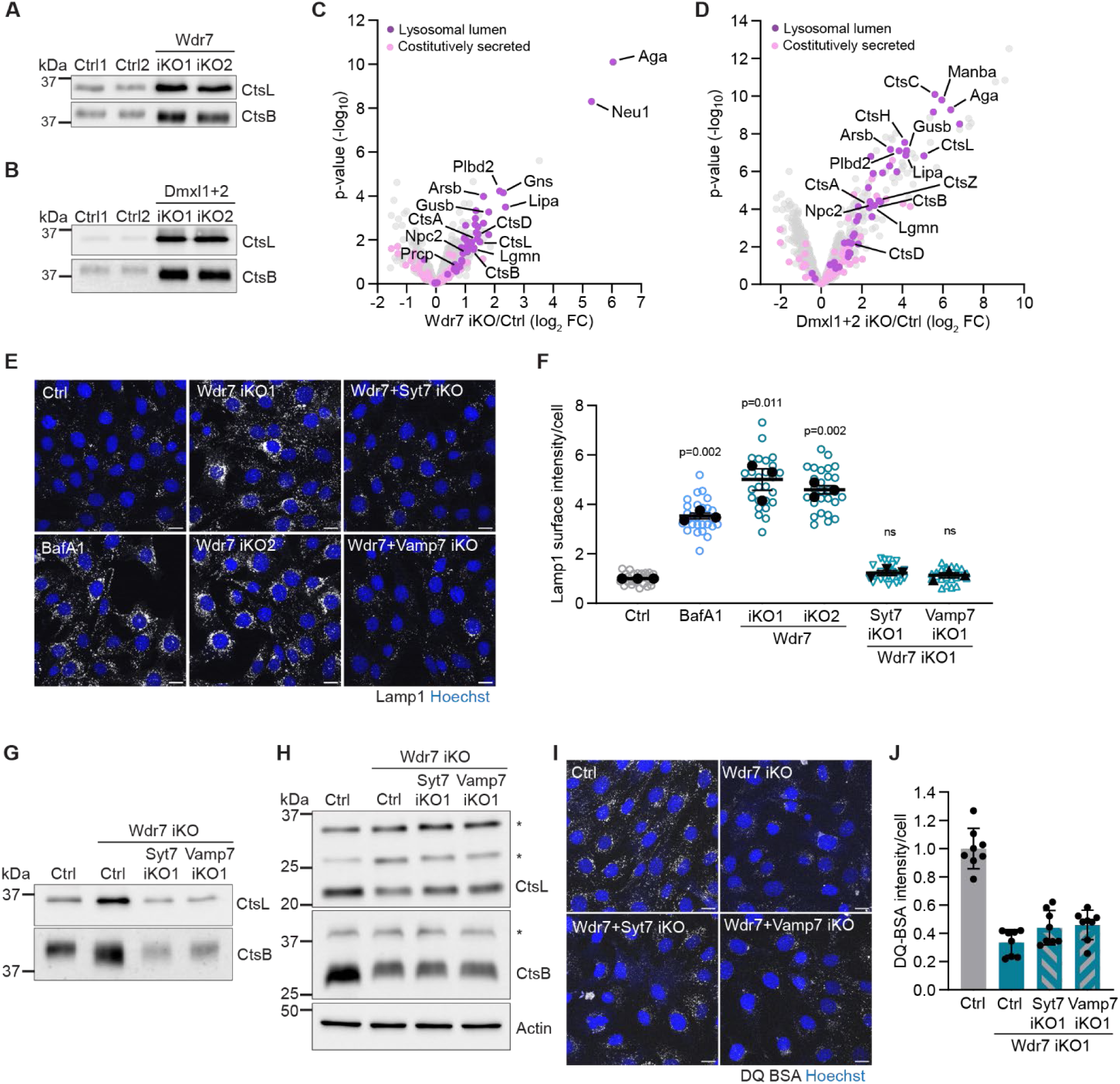
Loss of mRAVE results in lysosomal exocytosis. **(A**, **B)** Immature cathepsin secretion in Wdr7 iKO (A) and Dmxl1 + Dmxl2 iKO (B) MEFs, analyzed by immunoblotting. **(C, D)** Changes in luminal lysosomal proteins and constitutively secreted proteins in the secretome of Wdr7 iKO (C) and Dmxl1 + Dmxl2 iKO (D) MEFs, quantified by label-free LC– MS. **(E)** Lysosomal exocytosis in MEFs with Wdr7 iKO ± Syt7 or Vamp7 iKO or after 2 h bafilomycin A1 [1 µM], analyzed by IF of cell-surface Lamp1. **(F)** Quantification of cell-surface Lamp1 in cells as shown in (E). **(G)** Immature cathepsin secretion in Wdr7 iKO MEFs ± Syt7 iKO or Vamp7 iKO, analyzed by immunoblotting. **(H)** Mature cathepsin abundance in Wdr7 iKO MEFs ± Syt7 iKO or Vamp7 iKO, analyzed by immunoblotting. Asterisk denotes immature proenzyme. **(I)** DQ BSA degradation in Wdr7 iKO MEFs ± Syt7 iKO or Vamp7 iKO, analyzed by live-cell imaging. **(J)** Quantification of DQ BSA fluorescence in cells shown in (I). (**C, D)** n = 5 independent experiments. (**F**) Data are replicate mean ± SEM (closed circles; n = 3 independent experiments) and fields of view (open circles; ≥ 8 per replicate). (**J**) Data are mean ± SD and fields of view (n = 8). *p* values were calculated using a two-tailed unpaired t-test with Welch correction. Scale bars, 20 µm.

Perturbed lysosomal acidification can trigger lysosomal exocytosis, a process during which lysosomes fuse with the plasma membrane, resulting in cell surface exposure of lysosomal membrane proteins and release of luminal contents (Miao *et al*, 2015; Tancini *et al*, 2020). To determine whether exocytosis accounts for lysosomal enzyme secretion in mRAVE-deficient cells, we performed cell surface staining for the luminal domain of the lysosomal transmembrane protein Lamp1 (Reddy *et al*, 2001). Loss of Wdr7 led to a strong increase in Lamp1 at the cell surface, similar to the effect of inhibiting V-ATPases with bafilomycin A1 (Fig. 5E, F). To directly test the involvement of lysosomal exocytosis, we genetically ablated Syt7 or Vamp7, which are required for fusion of lysosomes with the plasma membrane (Rao *et al*, 2004). Co-depletion of Syt7 or Vamp7 together with Wdr7 suppressed the cell-surface localization of Lamp1 (Fig. 5E, F). Consistently, depleting Syt7 or Vamp7 in Wdr7-deficient cells abrogated the increased secretion of cathepsins B and L (Fig. 5G). Thus, secretion of immature lysosomal enzymes in mRAVE-deficient cells occurs through the lysosomal exocytosis pathway. Notably, suppressing lysosomal exocytosis in Wdr7-deficient cells did not restore intracellular levels of mature cathepsins (Fig. 5H) or rescue the defect in DQ BSA degradation (Fig. 5I, J). These results indicate that lysosomal exocytosis and the resulting secretion of lysosomal enzymes are a consequence of lysosomal dysfunction in mRAVE-deficient cells.

### mRAVE couples V-ATPase assembly to mTORC1-regulated control of lysosomal catabolism

The above results establish a central role for mRAVE in the assembly of V-ATPase complexes across multiple organelles. To determine which phenotypes depend specifically on the acidification of lysosomes, we sought to restore lysosomal pH independently of mRAVE. We ectopically expressed H⁺/Cl⁻ antiporters, ClC-6 and ClC-7, that localize to endolysosomal compartments (Fig. EV6A, B) and contribute to luminal acidification alongside the V-ATPase (Jentsch & Pusch, 2018; Settembre & Perera, 2024). Expression of ClC-6 restored lysosomal pH to control levels in Wdr7-deficient cells, while ClC-7 had a slightly weaker effect (Fig. 6A; Fig. EV6C). We therefore utilized ClC-6, whose expression in Wdr7-deficient cells did not rescue Golgi pH (Fig. EV6D) or V-ATPase assembly (Fig. EV6E) and restored lysosomal acidification in a proton transport-dependent manner (Fig. EV6F, G). Thus, ClC-6 expression provides a means to functionally uncouple lysosomal acidification from mRAVE-dependent V-ATPase assembly.

**Figure 6.**
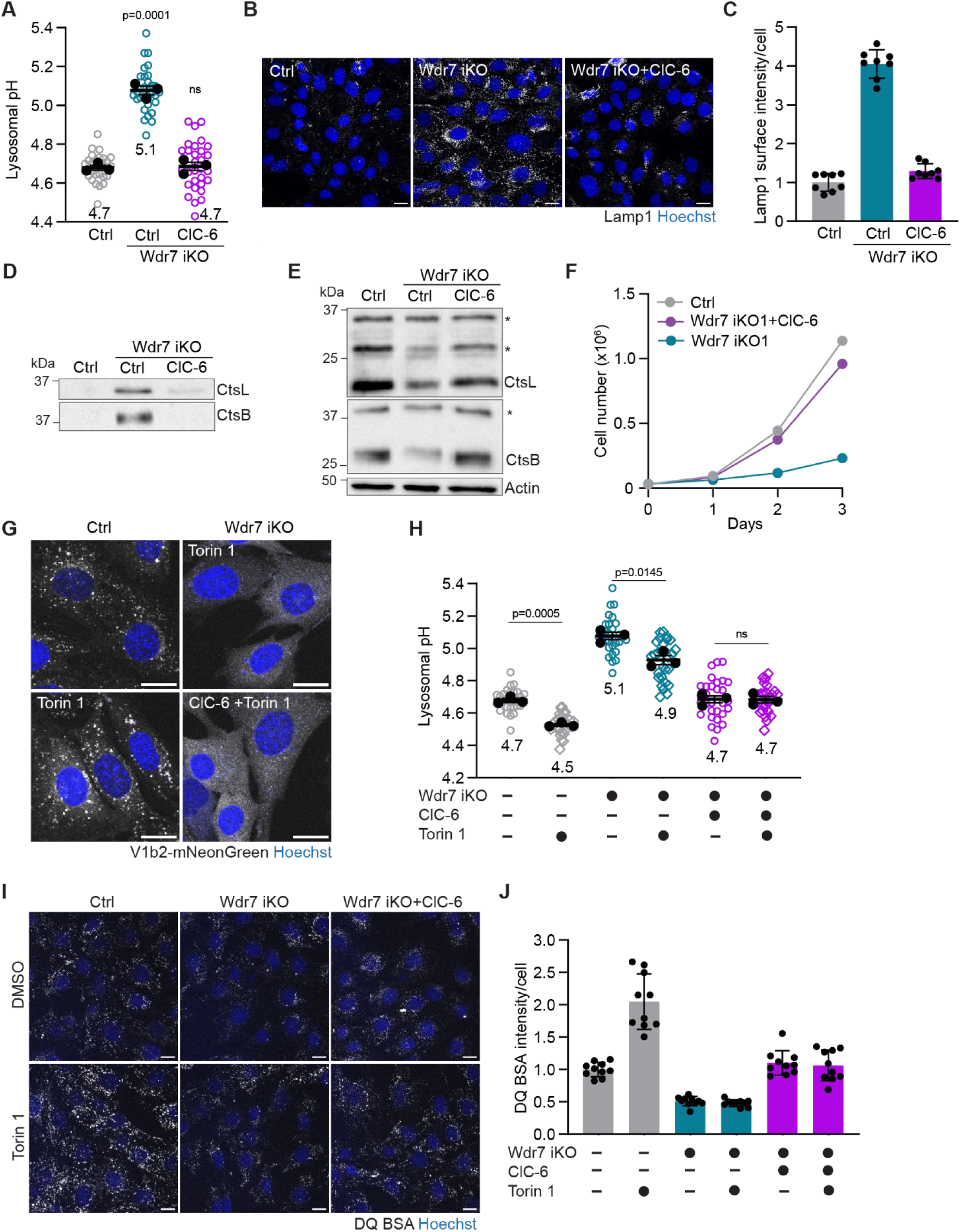
ClC-6 expression restores basal but not mTORC1-regulated lysosomal catabolism in the absence of mRAVE. **(A)** Lysosomal pH in Wdr7 iKO MEFs ± ClC-6 expression, quantified by ratiometric imaging of 10 kDa Dextran Oregon Green and Alexa Fluor 647. **(B)** Lysosomal exocytosis in Wdr7 iKO MEFs ± ClC-6 expression, analyzed by IF of cell-surface Lamp1. **(C)** Quantification of cell-surface Lamp1 in cells shown in (B). **(D)** Immature cathepsin secretion in Wdr7 iKO MEFs ± ClC-6 expression, analyzed by immunoblotting. **(E)** Mature cathepsin abundance in Wdr7 iKO MEFs ± ClC-6 expression, analyzed by immunoblotting. Asterisk denotes immature proenzymes. **(F)** Proliferation of Wdr7 iKO MEFs ± ClC-6 expression. **(G)** Subcellular localization of V1b2-mNeonGreen in Wdr7 iKO MEFs after 1 h ± torin 1 [400 nM], analyzed by live-cell imaging. **(H)** Lysosomal pH in Wdr7 iKO MEFs ± ClC-6 expression after 1 h ± torin 1 [400 nM], quantified by ratiometric imaging of 10 kDa Dextran Oregon Green and Alexa Fluor 647. **(I)** DQ BSA degradation in Wdr7 iKO MEFs ± ClC-6 expression after 1 h ± torin 1 [400 nM], analyzed by live-cell imaging. **(J)** Quantification of DQ BSA fluorescence in cells shown in (I). (**A, H**) Data are replicate mean ± SEM (closed circles; n = 3 independent experiments) and fields of view (open circles; ≥9 per replicate). (**C, J**) Data are mean ± SD and fields of view (n = 8–10). (**F**) Data are mean ± SD (n = 3 technical replicates). *p* values were calculated using a two-tailed unpaired t-test. Scale bar = 20 µm.

Restoring lysosomal acidification corrected multiple defects that arise from the loss of mRAVE: Expression of ClC-6 in Wdr7-deficient cells suppressed Lamp1 cell-surface localization (Fig. 6B, C) and prevented secretion of cathepsins B and L (Fig. 6D). Intracellularly, these changes were accompanied by restored levels of mature cathepsins, lysosomal proteolytic activity and lysotracker accumulation (Fig. 6E; Fig. EV6G–I). Whereas loss of Wdr7 severely compromised cell proliferation, Wdr7-deficient cells expressing ClC-6 proliferated at comparable rates as controls (Fig. 6F). These findings indicate that sustaining lysosomal acidification is the essential function of mRAVE in proliferating cells.

Finally, we examined the requirement for mRAVE in the adaptive regulation of lysosomal activity by mTORC1. In control cells, torin 1 treatment enhanced V-ATPase assembly, lysosomal acidification, and DQ BSA degradation (Fig. 6G–J). In contrast, Wdr7-deficient cells—despite restored basal acidification via ClC-6—failed to increase V-ATPase assembly, lysosomal acidification or lysosomal proteolysis upon torin 1 treatment (Fig. 6G–J). Thus, mRAVE is critical for V-ATPase assembly to enhance the acidification and catabolic activity of lysosomes in response to mTORC1 inactivation.

## Discussion

Reversible association of the V-ATPase V₁ and V₀ domains is a conserved mechanism to regulate organelle acidification. Here, we define mRAVE as the principal assembly factor for mammalian V-ATPases, which establishes luminal acidification across the endomembrane system, sustains lysosomal function and enables adaptive changes in lysosomal catabolism downstream of mTORC1 signaling.

The RAVE complex, composed of Rav1, Rav2 and Skp1, was identified in budding yeast as an organelle-specific V-ATPase assembly factor that promotes V_1_ – V_o_ domain association at the vacuole (Jaskolka *et al*., 2021b; Seol *et al*., 2001). This organelle selectivity results from the specific interaction of RAVE with the vacuolar Voa subunit, Vph1, but not with its Golgi-resident homologue, Stv1 (Smardon *et al*, 2014). By contrast, the present study demonstrates that mRAVE mediates the assembly of V-ATPase complexes containing the ubiquitously expressed subunits Voa1–3, which target proton pumps to lysosomes, endosomes and the Golgi in mammalian cells (Collins & Forgac, 2020; Nixon & Rubinsztein, 2024). Genetic studies in mice further indicate that mRAVE assembles kidney-specific Voa4-containing V-ATPases at the plasma membrane (Eaton *et al*., 2024). Thus, unlike its yeast counterpart, mRAVE is broadly required for V-ATPase assembly across different membrane compartments. Despite the evolutionary conservation of the V-ATPase, the metazoan RAVE-related complex differs substantially in size and composition (Lee *et al*., 2025; Wang *et al*, 2024; Winkley & Kane, 2025). The Dmxl1/2 subunit is more than twice as large as Rav1, and Wdr7 represents an additional large subunit absent from yeast RAVE. This expansion of the assembly factor may reflect the need to accommodate a broader diversity of V-ATPase complexes and integrate additional regulatory inputs in metazoan cells.

The regulation of V-ATPase assembly couples organelle acidification to stress responses and nutrient signaling. In yeast, RAVE restores acidification of the vacuole by promoting V-ATPase reassembly in response to glucose refeeding following starvation-induced disassembly (Jaskolka *et al*., 2021a; Smardon *et al*., 2002). In mammalian cells, mRAVE similarly restores acidification of lysosomes by promoting V-ATPase assembly in response to lysosomal membrane damage and resulting proton gradient collapse (Lee *et al*., 2025; Nardone *et al*., 2025). Beyond its function in organelle reacidification, our findings point to a broader physiological role of mRAVE. Under basal conditions, mRAVE is required to sustain V-ATPase levels and hence acidification of lysosomes and the Golgi, indicating ongoing assembly of proton pumps. In response to mTORC1 inactivation, mRAVE and V₁ are co-recruited to lysosomes to enhance V-ATPase assembly, luminal acidification and degradation of macromolecular contents. The mechanisms underlying RAVE recruitment to membranes remain incompletely understood, but direct interaction of yeast RAVE with the Vo domain may suffice for glucose-regulated recruitment to the vacuole (Jaskolka & Kane, 2020). In mammalian cells, lysosomal membrane damage has been shown to induce mRAVE recruitment to lysosomes via CASM (conjugation of ATG8 to single membranes), a non-canonical autophagy pathway (Durgan & Florey, 2022; Lee *et al*., 2025). mTORC1 is not involved in CASM, however, and how precisely it regulates lysosomal V-ATPase assembly remains unclear. Golgi V-ATPase are also assembled by mRAVE but do not respond to mTORC1, suggesting that regulation of V-ATPase assembly by nutrient signaling is specific to lysosomes.

Our findings demonstrate that the activity of lysosomes is remarkably sensitive to changes in luminal pH. In mRAVE-deficient cells, impaired V-ATPase assembly elevates lysosomal pH by ∼0.4, leading to the accumulation of dysfunctional lysosomes. mRAVE-deficient cells also show increased lysosomal exocytosis and secretion of luminal enzymes—reminiscent of the response to lysosomal deacidification during pathogen invasion (Miao *et al*., 2015)—yet lysosomal proteolysis remains largely suppressed even when exocytosis is blocked. This suggests that mRAVE-dependent V-ATPase assembly is critical to sustain a pH environment that supports cathepsin activation. Conversely, mTORC1 inactivation enhances the assembly of lysosomal V-ATPases, which decreases lysosomal pH by ∼0.3 and markedly boosts catabolic activity (the present study; (Ratto *et al*., 2022)). These functional consequences of subtle pH changes indicate that regulation of V-ATPase assembly enables cells to tune lysosomal acidification and, consequently, degradative activity. pH perturbations of similar magnitude have been observed in cancer, neurodegeneration and senescence (Eaton *et al*., 2021; Nixon & Rubinsztein, 2024; Tan & Finkel, 2023; Webb *et al*., 2021), suggesting that dysregulated lysosomal acidification could have broad pathological consequences.

## Acknowledgments

We are grateful to members of the Palm lab for experimental advice and helpful discussions. We thank V. Kalter and the DKFZ flow cytometry facility, the DKFZ light microscopy facility, D. Frey and the DKFZ proteomics facility. This work was supported by the Peter & Traudl Engelhorn Stiftung (A.Z.), the Health + Life Science Alliance Heidelberg Mannheim (A.Z.) and the International DKFZ PhD Program (N.S.S).

## Author contributions

N.S.S. and W.P. conceived the project and designed research together with A.Z. N.S.S. and A.Z. conducted research, with contributions from A.P. M.S. and D.H. conducted proteomics. All authors analyzed the data. N.S.S., A.Z. and W.P. wrote the manuscript, with input from all authors. W.P. supervised the study.

## Ethics declaration

The authors declare no competing interests.

## Data availability

The mass spectrometry data generated in this study have been deposited at ProteomeXchange via the PRIDE partner repository: Lyso-IP enrichment and Lyso-IP Wdr7 iKO (PXD070950), HA-V1b2 co-IP (PXD070848), cellular proteome Wdr7 iKO (PXD070900) and secretome Wdr7 iKO and Dmxl1 + Dmxl2 iKO (PXD070740).

## Methods

### Reagents and tools table

| Reagent/Resource | Reference or Source | Identifier or Catalog Number |
| --- | --- | --- |
| **Experimental Models** |  |  |
| iCas9 MEFs | Ratto et al. (Ratto *et al.*, 2022) | N/A |
| iCas9 EPP2 | Pechincha et al. (Pechincha *et al.*, 2022) | N/A |
| iCas9 MIA PaCa-2 | Pechincha et al. (Pechincha *et al.*, 2022) | N/A |
| iCas9 RKO | Groessl et al. (Groessl *et al.*, 2025) | N/A |
| Wild type MEFs | Ratto et al. (Ratto *et al.*, 2022) | N/A |
| HEK293T | ATCC | CRL-3216 |
| **Recombinant DNA** |  |  |
| pCMVR8.74 | Addgene | 22036 |
| pCMV-VSV-G | Addgene | 8454 |
| pRRL-SFFV-rtTA3-IRES-HygroR | Michlits et al. (Michlits *et al.*, 2020) | N/A |
| pRLL-TRE3G-Cas9-P2A-BFP | Michlits et al. (Michlits *et al.*, 2020) | N/A |
| Dual-hU6-sgRNA-mU6-sgRNA-Ef1a-Thy1.1-P2A-NeoR/-BlastR | Pechincha et al. (Pechincha *et al.*, 2022) | N/A |
| pMaCtag-Z07 | Addgene | 124790 |
| pMaCtag-Z11 | Addgene | 120054 |
| pRRL-UbC-HA-Atp6v1b2-mNeonGreen | This study | N/A |
| pRRL-UbC-HA-Atp6v1b2-mScarlet-I | This study | N/A |
| pLV-UbC-HA-Atp6v1b2 | This study | N/A |
| pLV-Ef1a-3xHA-Atp6v1b2-mNeonGreen | This study | N/A |
| pLV-UbC-3xFlag-Atp6v0a1-mScarlet-I | This study | N/A |
| pLV-UbC-3xFlag-Atp6v0a1 | This study | N/A |
| pLV-Ef1a-3xFlag-Atp6v0a2-mScarlet-I | This study | N/A |
| pLV-Ef1a-3xFlag-Atp6v0a2 | This study | N/A |
| pLV-UbC-3xFlag-Atp6v0a3-mScarlet-I | This study | N/A |
| pLV-UbC-3xFlag-Atp6v0a3 | This study | N/A |
| pLV-UbC-V5-Wdr7 | This study | N/A |
| pLV-UbC-HA-Wdr7-mNeonGreen | This study | N/A |
| pLV-UbC-V5-Wdr7 sgRNA-resistant | This study | N/A |
| pLV-UbC-CICN6-Flag | This study | N/A |
| pLV-UbC-CICN6<sup>E200A</sup>-Flag | This study | N/A |
| pLV-UbC-CICN7-Flag-P2A-Ostm1-V5 | This study | N/A |
| pLV-UbC-TMEM192-3xHA | This study | N/A |
| pLV-UbC-TMEM192-2xFlag | This study | N/A |
| MGAT-SEP-Ruby3 | Addgene | 200937 |
| **Antibodies** |  |  |
| rabbit anti-HA | Cell Signaling Technology | 3724 |
| rabbit anti-Golgin97 | Cell Signaling Technology | 13192 |
| rabbit anti-Lamp1 | Cell Signaling Technology | 3243 |
| rabbit anti-Rab7 | Cell Signaling Technology | 9367 |
| rabbit anti-Atp6v1b2 | Cell Signaling Technology | 14617 |
| rabbit anti-V5 | Cell Signaling Technology | 13202 |
| mouse anti-β-actin | Sigma-Aldrich | A5441 |
| goat anti-Cathepsin B | R&D Systems | AF965 |
| goat anti-Cathepsin L | R&D Systems | AF952 |
| mouse anti-Cathepsin D | R&D Systems | AF1029 |
| mouse anti-Flag M2 | Sigma-Aldrich | F1804 |
| rat anti-Lamp2 | Origene | AM08170BT-N |
| rabbit anti-Atp6v0d1 | Abcam | ab202899 |
| rabbit anti-Atp6v1a | Abcam | ab199326 |
| rabbit anti-Atp6v1e1 | ProteinTech | 15280-1-AP |
| rabbit anti-Lamp1 | Santa Cruz Biotechnology | sc-19992 |
| rabbit anti-CD63 | Abcam | ab217345 |
| rabbit anti-EEA1 | Cell Signaling Technology | 3288 |
| rabbit anti-Calreticulin | Cell Signaling Technology | 12238 |
| rabbit anti-TOM20 | ProteinTech | 11802-1-AP |
| rabbit anti-VDAC | Cell Signaling Technology | 4661 |
| rabbit anti-S6K | Cell Signaling Technology | 2708 |
| rabbit anti-Histone H3 | Cell Signaling Technology | 4499 |
| mouse anti-GM130 | BD Biosciences | 610822 |
| rabbit anti-Alexa Fluor 488 | Life Technologies | A11034 |
| rat anti-Alexa Fluor 488 | Life Technologies | A11006 |
| mouse anti-Alexa Fluor 568 | Life Technologies | A11004 |
| rat anti-Alexa Fluor 647 | Life Technologies | A21247 |
| HRP anti-goat | Life Technologies | 31402 |
| HRP anti-mouse | Cytiva | NA931 |
| HRP anti-rabbit | Cytiva | NA934 |
| HRP anti-rat | Jackson ImmunoResearch | 112-035-143 |
| **Oligonucleotides and other sequence-based reagents** |  |  |
| dsgRNA sequences | This study | Appendix Table 1 |
| cDNA cloning primers | This study | Appendix Table 2 |
| **Chemicals, Enzymes and other reagents** |  |  |
| Torin 1 | Tocris | 4247 |
| Nigericin | Tocris | 4312/10 |
| Bafilomycin A1 | Cayman | 11038 |
| Valinomycin | MedChemExpress | HY-N6693 |
| Monesin | Selleckchem | S2324 |
| Halt Protease and Phosphatase Inhibitor Cocktail | Thermo Fisher Scientific | 78444 |
| Doxycycline | Sigma-Aldrich | J67043 |
| PEI | Polysciences | 23966 |
| Blasticidin | Santa Cruz Biotechnology | sc-495389 |
| Hygromycin | Invitrogen | 10687010 |
| Geneticin | Invitrogen | 10131035 |
| Fatty acid-free BSA | EMD Millipore | 126579 |
| Clarity Max Western ECL substrates | Bio-Rad | 1705062 |
| SuperSignal West Atto | Thermo Fisher Scientific | A38554 |
| Triton X-100 | Sigma-Aldrich | X100 |
| Pierce anti-HA magnetic beads | Thermo Fisher Scientific | 88836 |
| Oregon Green 488 dextran 10 kDa | Life Technologies | D7170 |
| Alexa Fluor 647 dextran 10 kDa | Life Technologies | D22914 |
| FITC dextran 10 kDa | Life Technologies | D1820 |
| DQ Green BSA | Life Technologies | D12050 |
| Hoechst | Cayman | 15547 |
| LysoTracker Red | Life Technologies | L7528 |
| SiR-Lysosome | SpiroChrome | SC012 |
| Pierce BCA Protein Assay Kit | Fisher Scientific | 18064090 |
| Bio-Rad Protein Assay Kit | Bio-Rad | 5000001 |
| MycoAlert Mycoplasma Detection Kit | Lonza | 126579 |
| SuperScript IV Reverse Transcriptase | Thermo Fisher Scientific | 18090010 |
| Q5 High-Fidelity DNA Polymerase | NEB | M0491 |
| Q5 Site-Directed Mutagenesis Kit | NEB | E0554S |
| Gibson assembly mix | NEB | E2611S |
| QIAprep Spin Miniprep Kit | Qiagen | 27104 |
| DNeasy Blood & Tissue Kit | Qiagen | 69504 |
| QIAquick Gel Extraction Kit | Qiagen | 28704 |
| **Software** |  |  |
| NEBuilder Assembly Tool version 2.10.8 | NEB | N/A |
| ImageJ 1.54f (Fiji) | National Institutes of Health | N/A |
| Image lab 6.0.0.25 | Bio-Rad | N/A |
| Prism Version 10.4.1 | Graph Pad | N/A |
| ZEN Blue 3.7.97.07000 | Carl Zeiss Microscopy | N/A |
| ZEN Black 8.1.0.484 | Carl Zeiss Microscopy | N/A |
| Spectronaut 20.2.250922.92449 | Biognosys |  |
| MaxQuant 2.1.4.0 | Max Planck Institute of Biochemistry |  |
| R version 4.5.1 | R Foundation for Statistical Computing | N/A |
| R package limma_3.64.1 | Ritchie et al. (Ritchie *et al*, 2015) | N/A |
| R package vsn_3.76.0 | Huber et al. (Huber *et al*, 2002) | N/A |
| R package missForest_1.5 | Stekhoven et al.(Stekhoven & Bühlmann, 2012) | N/A |

### Cell lines and culture conditions

Experiments were conducted in SV40 large T antigen-immortalized embryonic fibroblasts (MEFs) (HA-V1b2 – V5-Wdr7 co-IP) and single-cell-derived doxycycline-inducible Cas9 (iCas9) cell lines (MEFs, EPP2, MIA PaCa-2 and RKO) that were described previously (Groessl *et al*., 2025; Pechincha *et al*., 2022; Ratto *et al*., 2022). Cells were cultured in DMEM/F-12 (Gibco #11320033) supplemented with 10% fetal bovine serum (Gibco #10270106) and 2 mM L-glutamine (Gibco #25030081). HEK 293T cells were used for virus production and cultured in DMEM (Gibco #2196903) supplemented with 10% FBS and 2 mM L-glutamine. All cell lines were cultured at 37 °C in a humidified incubator with 5 % CO₂.

Cell lines were routinely tested for mycoplasma contamination using the MycoAlertMycoplasma Detection Kit and authenticated by single nucleotide polymorphism (SNP) typing (Multiplexion) or by sequencing analysis. Cell culture reagents were obtained from Gibco unless otherwise specified.

### Lentivirus production and cell transduction

Lentiviral particles were produced by co-transfecting HEK 293T cells with the lentiviral expression vector, packaging plasmid pCMVR8.74 and envelope plasmid pCMV-VSV-G using polyethylenimine (PEI MW 25,000). Viral supernatants were collected 48 h post-transfection, filtered through a 0.45 μm PES filter and used to transduce target cells in the presence of 10 μg/mL polybrene (Sigma-Aldrich, #TR-1003) in a dilution of 1:10.

### Generation of gene knockouts

For inducible knockout (iKO) generation, iCas9 cell lines were transduced with lentiviral vectors encoding dsgRNAs targeting the respective gene: Dual-hU6-sgRNA-mU6-sgRNA-Ef1a-Thy1.1-P2A-NeoR, Dual-hU6-sgRNA-mU6-sgRNA-Ef1a-Thy1.1-P2A-BlastR or Dual-hU6-sgRNA-mU6-sgRNA-EF1αs-BFP. DsgRNA-expressing cells were selected with neomycin or blasticidin or sorted by FACS for BFP expression. Cells were cultured in antibiotics until seeded for the experiment to maintain dsgRNA expression, and Cas9 was induced with 0.6 μg/mL doxycycline for 4 d. Control dsgRNAs were Ctrl1 targeting a non-coding chromosome region (Chr1.1) and Ctrl2 targeting a non-expressed gene (CD19). dsgRNA sequences are listed in Table 1.

### Generation of cDNA expression constructs

Atp6v1b2, Atp6v0a1, Atp6v0a2, Atp6v0a3 and Wdr7 cDNAs were amplified by PCR from a mouse cDNA library generated by reverse transcription of mRNA isolated from MEFs; OSTM1 cDNA from a human cDNA library generated from RKO cells (human); ClCN6 (ClC-6) and ClCN7 (ClC-7) cDNAs from human Gateway ORF clones (provided by the Genomics and Proteomics Core Facility, DKFZ); and TMEM192-3xHA and TMEM192-2xFlag from Addgene plasmids #102930, #102929 (Abu-Remaileh *et al*., 2017). N- or C-terminal HA, Flag and V5 tags were introduced with the PCR primer. mNeonGreen and mScarlet-I were amplified from pMaCTag-Z07 (Addgene #124790) and pMaCTag-Z11 (Addgene #120054), respectively and fused to the cDNA of interest with a GGGS linker. CLCN7 and OSTM1 cDNAs were cloned into the same expression vector, separated by a P2A cleavage site, to express equal quantities of the two transporter subunits. sgRNA-resistant Wdr7 cDNA for genetic rescue experiments and proton transport-deficient ClC-6^E200A^ (Neagoe *et al*, 2010) were generated using the Qiagen Site-Directed Mutagenesis Kit.

Cloning primers were designed using the NEBuilder Assembly Tool, and cloning was performed with the Q5 HiFi polymerase and the HiFi DNA Assembly Master Mix following the provider’s protocols. Expression plasmids generated in this study are listed in the Key Resources Table. cDNA cloning primers are listed in Table 2.

### Cell proliferation assay

Cells were maintained in the appropriate antibiotic-containing medium, and Cas9 expression was induced with 0.6 µg/ml doxycycline for 4 d before the experiment. Cells were seeded in standard medium in 12-well plates in 3 technical replicates. Cell numbers were counted with a CASY Cell Counter (OMNI Life Science) at day 0 (at least 4 h after seeding to allow cell attachment) and then daily or on day 3. Fold-changes in cell number from day 0 to day 3 were calculated as indicated.

### Immunoblotting

Cells were washed with ice-cold PBS and lysed in lysis buffer (50 mM HEPES pH 7.4, 40 mM NaCl, 2 mM EDTA, 1% Triton X-100, supplemented with 1 mM sodium orthovanadate, 50 mM sodium fluoride, 10 mM sodium pyrophosphate, 10 mM sodium glycerophosphate and Halt protease and phosphatase inhibitor cocktail). Lysates were incubated for 15 min on ice and cleared by centrifugation at 18,000 × g for 5 min at 4 °C. Protein concentration was measured by BCA assay.

To analyze secreted proteins, cells were cultured in Opti-MEM for 24 h. Cellular supernatants were collected centrifuged at 1,000 × g for 5 min, followed by centrifugation at 18,000 × g for 20 min at 4°C. Cell numbers were quantified on parallel culture plates with a CASY Cell Counter and used to normalize the volume of cellular supernatants.

Proteins were denatured in Laemmli buffer containing 10 % β-mercaptoethanol for 10 min at 95 °C. Equal amounts of protein were analyzed by SDS gel electrophoresis and immunoblotting on nitrocellulose membranes following standard protocols. When samples were probed across multiple membranes, blotting was performed in parallel under identical conditions. Primary antibodies were used at a 1:1,000 dilution in TBS-T (20 mM Tris, 150 mM NaCl and 0.1 % Tween) with 5 % BSA, secondary antibodies at a 1:5,000 dilution in TBS-T with 5 % skim milk. Immunoblots were imaged with the ChemiDoc Touch system (Bio-Rad) using ECL substrates Clarity Max Western. For stripping, membranes were incubated in stripping buffer (200 mM glycine, 0.1 % SDS, pH 2.2), washed three times with TBST (10 min each) and re-blocked in 5 % milk in TBST for 30 min. Band integrated intensities were quantified using Fiji.

### Isolation of lysosome-enriched membrane fractions

Lysosomes were loaded overnight with 2 mg/ml 10 kDa dextran (Sigma-Aldrich), followed by a 4 h chase in fresh medium. Cells were washed, resuspended in homogenization buffer (20 mM HEPES pH 7.5, 125 mM KCl, 50 mM sucrose, 1 mM EDTA, protease and phosphatase inhibitors) and homogenized using 12 strokes of a KIMBLE Dounce tissue grinder (large clearance pestle, Sigma-Aldrich). Post-nuclear supernatants were obtained by centrifugation at 800 × g for 5 min at 4 °C. Membranes were subsequently pelleted at 18,000 × g for 20 min at 4 °C. Pellets were lysed as above and analyzed by immunoblotting. Protein concentrations were determined by Bradford assay.

### Co-immunoprecipitation

For co-IP, cells were seeded for immunoblotting (3×10^5^ cells in 6-well plates) or proteomics (4×10^6^ cells in 15 cm dishes) 1 day before the experiment. Cells were washed 1x with ice-cold PBS, scraped with 300 µL (6-well plates) or 1 mL (15-cm dishes) of IP buffer (V_1_ – V_o_ co-IP: 40 mM HEPES pH 7.4, 150 mM NaCl, 2 mM EDTA, 2.5 mM MgCl₂, 5% glycerol; V_1_ – Wdr7 co-IP: 40 mM HEPES pH 7.4, 150 mM NaCl; 1 % Triton X-100, protease and phosphatase inhibitors were added right before use), collected into pre-cooled tubes and incubated on ice for 15 min. Lysates were centrifuged at 11,500 rpm for 5 min at 4 °C and supernatants transferred to new pre-cooled tubes. During lysis, magnetic beads were equilibrated by washing 3x with IP buffer. 10 % of the lysate was collected as input and mixed with 4x Laemmli buffer. The remaining lysate was incubated with antibody-coupled magnetic beads for 2 h at 4 °C with end-to-end rotation. Beads were washed 1x with IP buffer, resuspended in 2x Laemmli and heated in a thermocycler at 65 °C for 10 min at 500 rpm. Beads were removed using a magnetic rack and supernatants used for SDS–PAGE or LC–MS analysis. 1 – 2 % of input samples were loaded as indicated.

### Lysosomal immunoprecipitation

For lysosomal immunoprecipitation (Lyso-IP), 3 × 10^6^ MEFs stably expressing pLV-UbC-TMEM192-3xHA (lyso-IP) or pLV-UbC-TMEM192-2xFlag (control-IP) were seeded into 15 cm dishes one day before the experiment. Magnetic beads were equilibrated by washing 3x times with 1 mL KPBS (136 mM KCl, 10 mM KH₂PO₄, pH 7.25, protease and phosphatase inhibitors) and kept on ice. Cells were washed twice with ice-cold PBS, scraped into 1 mL ice-cold KPBS, centrifuged at 1,000 × g for 2 min at 4 °C and resuspended in 950 µL KPBS. 50 µL of cell suspension were taken as input sample. Remaining suspensions were homogenized with 20 strokes in a KIMBLE Dounce tissue grinder (large clearance pestle, Sigma-Aldrich) on ice. Homogenates were centrifuged at 1,000 × g for 2 min at 4 °C. and incubated with 60 µL of equilibrated magnetic beads for 5 min at 4 °C with end-to-end rotation. Beads were washed 3x with ice-cold KPBS, resuspended in 90 µL lysis buffer and incubated for 10 min on ice. Proteins were eluted from beads and denatured by incubation with 4x Laemmli buffer at 65 °C for 15 min. Input samples were incubated with 90 µL lysis buffer for 20 min on ice, centrifuged at 13,000 × g for 10 min at 4 °C and resulting supernatants mixed with 4x Laemmli buffer.

### Sample preparation for proteomics

For Lyso-IP and co-IP analyses, samples were prepared as described above. For cellular proteome analysis, cells were washed twice with ice-cold DPBS, scraped into 1 mL ice-cold DPBS and centrifuged at 1,000 × g for 2 min at 4 °C. Pellets were resuspended in lysis buffer (50 mM HEPES pH 7.4, 40 mM NaCl, 2 mM EDTA, 1% Triton X-100, supplemented with 1 mM sodium orthovanadate, 50 mM sodium fluoride, 10 mM sodium pyrophosphate, 10 mM sodium glycerophosphate and Halt protease and phosphatase inhibitor cocktail (Thermo Fisher), incubated for 20 min on ice and cleared by centrifugation at 13,000 × g for 10 min at 4 °C. Supernatants were collected and mixed with 4x Laemmli buffer. For all analyses 5 µg of protein were digested with trypsin with an AssayMAP Bravo liquid handling system (Agilent technologies) running the autoSP3 protocol (Müller *et al*, 2020).

For secretome analysis, cellular supernatants were collected as described above, centrifuged for 5 min at 15,000 rpm at room temperature and transferred in fresh tubes. Samples were mixed with an equal amount of RIPA buffer, and protein concentration was determined by BCA assay. 10 µg of proteins were digested with trypsin with an AssayMAP Bravo liquid handling system (Agilent technologies) running the autoSP3 protocol (Müller *et al*., 2020).

### LC-MS/MS analysis

Dried peptide samples were reconstituted (97.4% H_2_O, 2.5% hexafluoro-2-propanol and 0.1% trifluoroacetic acid), and one third was used for subsequent analysis. LC– MS/MS was carried out using a Vanquish Neo UPLC system (Thermo Fisher Scientific) for Lyso-IP, cell lysate and secretome analyses or an Ultimate 3000 UPLC system (Thermo Fisher Scientific) for co-IP analysis, directly connected to an Orbitrap Exploris 480 mass spectrometer for a total of 60 min (Lyso-IP, secretome), 90 min (co-IP) or 120 min (cell lysate) per sample. Peptides were online desalted on a trapping cartridge (Acclaim PepMap300 C18, 5 µm, 300 Å wide pore; Thermo Fisher Scientific) with a loading volume of 60 µl using 30 µl/min flow of 0.05 % trifluoroacetic acid in H_2_O. The analytical multistep gradient (300 nl/min) was performed with a nanoEase MZ Peptide analytical column (300 Å, 1.7 µm, 75 µm x 200 mm, Waters) using solvent A (0.1% formic acid in H_2_O) and solvent B (0.1% formic acid in acetonitrile). For 45 min (secretome, Lyso-IP), 72 min (co-IP) or 104 min (cell lysate) the concentration of B was linearly ramped from 4 % (cell lysate, co-IP) or 5 % (secretome, Lyso-IP) to 30 %, followed by a quick ramp to 78 % (co-IP) or 80 % (secretome, cell lysate, Lyso-IP), after 2 min (co-IP) or 4 min (secretome, cell lysate and Lyso-IP) the concentration of B was lowered to 2 % and a 10 min (co-IP) or a three column volumes (secretome, cell lysate, Lyso-IP) equilibration appended. Eluting peptides were analyzed in the mass spectrometer using data independent acquisition (DIA) mode (secretome, cell lysate, Lyso-IP) or data dependent acquisition (DDA) mode (co-IP).

For DIA, a full scan at 120k resolution (350-1400 m/z, 300 % AGC target, 45 ms maxIT) was following 20 (secretome), 25 (Lyso-IP) or 40 (cell lysate) DIA windows. The DIA acquisition covered a mass range of 400-1000 m/z (secretome, cell lysate) or 380-880 m/z (Lyso-IP) using windows of a variable width. Windows overlapped by 1 m/z, the AGC target was 1000 % with a maxIT of 54 ms and spectra were recorded at a resolution of 30k.

For DDA, a full scan at 60k resolution (380-1400 m/z, 300 % AGC target, 45 ms maxIT) was followed by up to 2 s of MS/MS scans. Peptide features were isolated with a window of 1.4 m/z, fragmented using 26 % NCE. Fragment spectra were recorded at 15k resolution (100 % AGC target, 54 ms maxIT). Unassigned and singly charged eluting features were excluded from fragmentation and dynamic exclusion was set to 30 s.

Each sample was followed by a wash run (22 min) to minimize carry-over between samples. Instrument performance throughout the course of the measurement was monitored by regular (approx. one per 48 h) injections of a standard sample and an in-house shiny application.

### Proteomics data analysis

Analysis of DIA RAW files was performed with Spectronaut (Biognosys, version 20.2.250922.92449 (Bruderer *et al*, 2015)) in directDIA+ (deep) library-free mode. Default settings were applied with the following adaptions. Within DIA Analysis under Identification, the Precursor PEP Cutoff was set to 0.01, the Protein Qvalue Cutoff (Run) to 0.01, the Protein PEP Cutoff to 0.01 and Run-level Protein scoring to Highest Scoring Observation (SN19 default). In Quantification, the Proteotypicity Filter was set to Only Protein Group Specific and the Protein LFQ Method to MaxLFQ, Cross-Run Normalization was disabled (secretome, Lyso-IP) or stayed enabled (cell lysate) and the Quantification window was set to Not Synchronized (SN17). In Post Analysis Use All MS-Level Quantities was set to True. The data was searched against the mouse proteome from Uniprot (mouse reference database with one protein sequence per gene, containing 21,757 unique entries from January 2025), the Cas9 sequence (secretome, cell lysate, Lyso-IP) and the contaminants FASTA from MaxQuant (246 unique entries from 22^nd^ of August 2025). Conditions were included in the setup.

For DDA, data analysis was carried out in MaxQuant (version 2.1.4.0, Tyanova et al. (Tyanova *et al*, 2016)), using an organism specific database extracted from Uniprot (mouse reference database with one protein sequence per gene, containing 21,757 unique entries from January 2025). Default settings were applied with the following adaptions. Match between runs (MBR) was enabled to transfer peptide identifications across RAW files based on accurate retention time and m/z. Fractions were set in a way that MBR was only performed within replicates. Label-free quantification (LFQ) was enabled with default settings. The iBAQ-value (Schwanhäusser *et al*, 2011) generation was enabled.

### Proteomics statistical analysis

For the statistical analysis, log2-transformed quantity values (secretome, cell lysate, Lyso-IP) or intensity values (co-IP) were used. The following steps were performed per statistical contrast and not across the whole data matrix. Protein groups with valid values in 70 % of the samples of at least one condition were used for statistics. Variance stabilization normalization (Huber *et al*., 2002) was applied to normalize across samples. Adapted from the Perseus recommendations (Tyanova & Cox, 2018), missing values being completely absent in one condition were imputed with random values drawn from a downshifted (2.2 standard deviations) and narrowed (0.3 standard deviations) intensity distribution of the individual samples. For missing values with no complete absence in one condition the R package missForest was used for imputation (Stekhoven & Bühlmann, 2012). The statistical analysis was performed with the R-package limma (Ritchie *et al*., 2015); single contrasts were adapted from the user guide chapter *Two Group*. The setup was adapted for proteomics data via setting the *eBayes* options *trend* and *robust* to *TRUE*. Proteins with a minimum of 2 identified peptides in at least 3 replicates were considered for further analysis. Wdr7 iKO Lyso-IP was analyzed using a subproteome curated from lysosome-enriched proteins identified in this study, the UniProt reference proteome ‘mouse lysosome and (Ratto *et al*., 2022; Wyant *et al*, 2018).

### Live-cell imaging

Live-cell imaging was performed in a humidified chamber at 37 °C and 5 % CO₂. Hoechst 33342 (0.5 µg/mL) was added 30 min before imaging. Where indicated, cells were treated with torin 1 [400 nM] directly on the microscope stage. Confocal microscopy was performed using a Zeiss LSM780 SD or LSM900 Airyscan 2 microscope equipped with a 40x or 63x (NA 1.40) oil immersion objective.

For live imaging, cells were seeded in 8-well chambered coverslips (IBIDI) and left to attach overnight. Lysosomal proteolysis, abundance and acidity were monitored by incubating cells with 0.1 mg/mL DQ Green BSA for 4 h and 50 nM LysoTracker Red for 1 h prior to imaging. Activated cathepsin D was measured by incubating cells with 1 µM SiR Lysosome for 1 h. V-ATPase-dependent lysosomal re-acidification was measured by FITC-dextran quenching. MEFs were loaded overnight with 1 mg/mL FITC dextran (10 kDa), followed by a 4 h chase. V-ATPase-mediated re-acidification of lysosomes was monitored by quantifying the rate of FITC fluorescence quenching in neutralized lysosomes after relief of V-ATPase inhibition. To this end, cells were treated with 20 nM bafilomycin A1 for 30 min, bafilomycin A1 was then removed by washing cells with medium containing 5 mg/mL fatty acid-free BSA, and FITC fluorescence quenching was imaged over time.

### Organelle pH measurements

To measure lysosomal pH, cells were seeded in 8-well chambered coverslips (IBIDI). On the next day, cells were loaded with 0.1 mg/mL Oregon Green 488 dextran (10 kDa) and 0.1 mg/mL Alexa Fluor 647 dextran (10 kDa) for 4 h. After a 4 h chase in dye-free medium, cells were treated as indicated. To generate a pH calibration curve, cells were incubated with calibration buffers (125 mM KCl, 25 mM NaCl, 25 mM HEPES/MES, pH 4.0–6.5) containing 10 µM nigericin, 10 µM monensin and 10 µM valinomycin to clamp lysosomal pH to the buffer. Fluorescence was measured by confocal microscopy and lysosomal pH calculated based on the ratio of Oregon Green (pH-sensitive) to Alexa Fluor 647 (pH-insensitive) fluorescence using the standard curve.

To measure Golgi pH, cells were stably transduced with MGAT-SEP-mRuby3 (Liu *et al*., 2023) and sorted by flow cytometry using 488 nm and 568 nm lasers to establish a cell population with homogeneous expression of both fluorescent proteins (50–70% fluorescence brightness). For imaging, cells were seeded in 8-well chambered coverslips (IBIDI). The following day, cells were stained with Hoechst 30 min prior to imaging. To generate a pH calibration curve, control cells were incubated in calibration buffers (125 mM KCl, 25 mM NaCl, 25 mM HEPES/MES, pH 6.0–7.5) containing 10 µM nigericin, 10 µM monensin and 10 µM valinomycin to clamp intracellular pH to the buffer. Fluorescence was measured by confocal microscopy and Golgi pH calculated based on the ratio of SEP (pH-sensitive) to mRuby3 (pH-insensitive) fluorescence using the standard curve.

### Immunofluorescence staining

Cells were seeded onto IBIDI chamber slides and cultured for 16–24 h. Cells were then rinsed with ice-cold PBS, fixed with 4 % paraformaldehyde in PBS for 15 min at room temperature and permeabilized with 0.05 % Triton X-100 in PBS for 5 min. After washing with PBS, cells were blocked for 30 min in blocking solution (4 % normal goat serum, 0.5 % BSA in PBS). Primary antibodies were diluted 1:400 in blocking solution and applied for 2 h at room temperature. After two washes with blocking solution, cells were incubated with Alexa Fluor-conjugated secondary antibodies (1:1,000 dilution) in blocking solution for 1 h. Cells were washed with PBS, stained with 10 μg/mL Hoechst 33342 in PBS for 5 min, washed twice with PBS and imaged directly. For cell-surface staining of LAMP1, cells were stained as above but the permeabilization step was omitted.

### Microscopy image analysis

For microscopy image analysis, fluorescence intensity was quantified using the particle analyzer function of Fiji (Schindelin *et al*, 2012) in randomly chosen fields of view across the entirety of each sample. Mean cellular fluorescence was determined by normalizing the integrated signal density of the respective fluorescent probe to cell number. Pearson’s coefficients were calculated using the JACoP plugin in Fiji. Where indicated, individual data points in super blots were normalized to the mean value of the corresponding control group.

### Statistical analysis

Data are represented as mean ± SD for technical replicates and independent biological replicates with single observation or as mean ± SEM for independent biological experiments with multiple observations. Statistical analyses were performed using Excel for immunoblot quantifications and Graphpad Prism for all the other quantifications. *p* values were calculated using a two-tailed unpaired t-test assuming equal variance; for data in which controls were normalized to 1, *p* values were calculated using a two-tailed unpaired t-test with Welch correction.

## Supplementary Information

**Expanded View** Figures EV1 – EV6

**Appendix** Appendix Tables 1 and 2

**Dataset EV1 Proteomics Lyso-IP enrichment (related to Figure 1**). Quantitative proteomics of lysosome-enriched fraction from MEFs expressing Tmem192-3xHA (Lyso-IP) or Tmem192-2xFlag (Ctrl-IP) via HA immunoprecipitation. Log2 fold-changes, *p* values and replicate precursor peptides for Lyso-IP compared to Ctrl-IP are provided.

**Dataset EV2 Proteomics Lyso-IP Wdr7 iKO (related to Figure 1**). Quantitative proteomics of lysosome-enriched fraction from MEFs expressing Tmem192-3xHA via HA immunoprecipitation. Log2 fold-changes, *p* values and replicate precursor peptides for Wdr7 iKO compared to Ctrl are provided. Ctrl is the same Lyso-IP sample as in Table S1. Lysosomal proteins shown in Fig. 1G are additionally listed separately.

**Dataset EV3 Proteomics HA-V1b2 co-IP (related to Figure 3**). Quantitative proteomics of HA co-IP from MEFs expressing HA-V1b2. Log2 fold-changes, *p* values and replicate unique peptides for HA-V1b2 compared to empty vector (Ctrl) are provided.

**Dataset EV4 Cellular proteome Wdr7 iKO (related to Figure 4**). Quantitative proteomics of cell lysates from MEFs. Log2 fold-changes, *p* values and replicate precursor peptides for Wdr7 iKO compared to Ctrl are provided.

**Dataset EV5 Secretome Wdr7 iKO and Dmxl1 + Dmxl2 iKO (related to Figure 5**). Quantitative proteomics of cellular supernatants from MEFs. Log2 fold-changes, *p* values and replicate precursor peptides for Wdr7 iKO or Dmxl1 + Dmxl2 iKO compared to Ctrl are provided.

